# A Germline Point Mutation in the MYC-FBW7 Phosphodegron Initiates Hematopoietic Malignancies

**DOI:** 10.1101/2023.10.23.563660

**Authors:** Brian Freie, Patrick A. Carroll, Barbara J. Varnum-Finney, Vijay Ramani, Irwin Bernstein, Robert N. Eisenman

**Affiliations:** Basic Sciences Division, Fred Hutchinson Cancer Center, Seattle WA, USA; Clinical Research Division, Fred Hutchinson Cancer Research Center, Seattle WA, USA; Gladstone Institute for Data Science and Biotechnology, University of California, San Francisco, San Francisco CA, USA

## Abstract

Oncogenic activation of MYC in cancers predominantly involves increased transcription rather than coding region mutations. However, MYC-dependent lymphomas frequently contain point mutations in the MYC phospho-degron, including at threonine-58 (T58), where phosphorylation permits binding by the FBW7 ubiquitin ligase triggering MYC degradation. To understand how T58 phosphorylation functions in normal cell physiology, we introduced an alanine mutation at T58 (T58A) into the endogenous *c-Myc* locus in the mouse germline. While MYC-T58A mice develop normally, lymphomas and myeloid leukemias emerge in ∼60% of adult homozygous T58A mice. We find that primitive hematopoietic progenitor cells from MYC-T58A mice exhibit aberrant self-renewal normally associated with hematopoietic stem cells (HSCs) and upregulate a subset of Myc target genes important in maintaining stem/progenitor cell balance. Genomic occupancy by MYC-T58A was increased at all promoters, compared to WT MYC, while genes differentially expressed in a T58A-dependent manner were significantly more proximal to MYC-bound enhancers. MYC-T58A lymphocyte progenitors exhibited metabolic alterations and decreased activation of inflammatory and apoptotic pathways. Our data demonstrate that a single point mutation in Myc is sufficient to produce a profound gain of function in multipotential hematopoietic progenitors associated with self-renewal and initiation of lymphomas and leukemias.

## INTRODUCTION

The *MYC* gene family is essential for normal development and cell physiology and is de-regulated in an extraordinarily wide range of cancers. The c-*MYC* gene (herein referred to as *MYC*) is induced in response to numerous cytokines, growth factors or other mitogenic signals. Serum and ligand stimulated pathways (such as Notch, Sonic Hedgehog, Wnt, EGF, PDGF, and others) activate *MYC* gene expression as an early and robust target. However, dependent on context, this response can be limited by the tight regulation of MYC at multiple levels (e.g., transcription initiation and pausing, RNA stability and export, translation, and protein half-life, (for recent review see (Dhanasekaran et al. 2022)).

In a large subset of cancers, the tight regulation of MYC is lost; *MYC* RNA is increased and is not down-regulated in response to differentiation signals or mitogen removal, resulting in increased protein abundance. This deregulation of *MYC* expression can be due to rearrangements involving *MYC* genomic loci, including chromosomal translocations, gene amplification, viral insertions and transductions, or gain-of-function mutations in one or more signal transduction pathways known to induce *Myc* expression (Dalla-Favera et al. 1982; Nowell et al. 1983; Adams et al. 1985; He et al. 1998). In each case, *MYC* expression is uncoupled from its normal physiological regulators. Indeed, even relatively low levels of de-regulated *MYC* expression can cause profound changes in cell biology (Murphy et al. 2008).

MYC activity is also regulated by modification of MYC protein. Perhaps the most crucial region of the MYC protein required for this post-translational regulation is located in the transactivation domain in a conserved region known as MYC Box I (for review see (Farrell and Sears 2014)). The established function of this region is as a phospho-degron, in which phosphorylation of MYC contributes to the regulation of MYC protein stability. Within murine MYC Box 1, the amino acid sequence from residues 57-63 (PTPPLSP) (corresponding to resides 72-78 in human MYC) are of critical importance. The growth-regulated kinase GSK3β phosphorylates threonine at amino acid 58 (T58) resulting in binding by the F-box protein Fbw7 (a subunit of the SCF ubiquitin ligase complex), which ubiquitinates MYC, targeting it for proteasome-mediated degradation (Gregory et al. 2003; Welcker et al. 2004; Yada et al. 2004). A recent study shows that MYC-FBW7 interaction involves an FBW7 dimer and is also stabilized by phosphorylation at MYC T244 (Welcker et al. 2022). MYC protein containing non-phosphorylated T58 is not efficiently bound by Fbw7 and exhibits increased stability. Mitogen-induced cell proliferation also triggers phosphorylation of S62 within MYC Box 1 through RAS-ERK signaling and CDKs. Phosphorylation of S62 primes GSK3β binding to, and subsequent phosphorylation of, T58 (Sears et al. 2000). Thus, MYC degradation is controlled through multiple kinases and closely tied to signal transduction pathways.

Early studies found that point mutations of T58, as adjacent amino acid residues in MYC Box I, that disrupt T58 phosphorylation were often observed in translocated *MYC* alleles in Burkitt’s lymphoma, AIDS related lymphomas, and B cell acute lymphoblastic lymphoma (Bhatia et al. 1993; Bhatia et al. 1994). Furthermore, three of the four transforming retroviruses from which the v-myc oncogene was originally identified also contain mutations in the cognate MYC Box I threonine (Bhatia et al. 1993). Therefore, mutations that disrupt T58 phosphorylation are a selected genetic event in MYC associated cancers. When MYC mutated at threonine 58 (T58A) was ectopically overexpressed by retrovirus in mouse hematopoietic cells, B cell lymphomas evolved faster than those generated by overexpressing normal, wild-type MYC (Hemann et al. 2005). In other studies, MYC T58A overexpressed in mouse mammary tissue induced mammary carcinomas more rapidly than wild-type c-MYC (Wang et al. 2011). These studies indicated that MYC blocked for phosphorylation at T58 is associated with increased oncogenicity compared to wild-type MYC. Indeed, T58 mutated MYC has been widely used in experimental models to increase and accelerate tumorigenesis (Swartling et al. 2012; Chalishazar et al. 2019). Moreover, ectopic expression of MYC-T58A maintained pluripotency in murine ES cells independent of LIF (Cartwright et al. 2005).

Interestingly, only a partial increase in MYC protein stability is seen when T58 is mutated, consistent with the fact that MYC stability is controlled by other pathways independent of T58 phosphorylation (Kim et al. 2003; Welcker et al. 2004; Schukur et al. 2020). Furthermore, evidence suggests that the T58 mutation is frequently acquired in tumors that already contain deregulated, and highly expressed, MYC (Bhatia et al. 1994). In addition, while it has been well established that deregulated MYC triggers apoptosis, the T58A mutant MYC actually results in decreased apoptosis when overexpressed (Hemann et al. 2005). Taken together, these findings suggest that T58 phosphorylation may affect MYC activity by coupling degradation with MYC function in an as yet undefined way.

The previous studies on T58A and other MYC mutations were carried out using exogenous, overexpressed, and deregulated MYC. To separate the effects of T58 mutation from the profound effects observed when MYC is overexpressed, we sought to study MYC regulation by T58 phosphorylation in a setting in which MYC expression is not overtly deregulated. To do this, we introduced a point mutation in murine MYC by changing T58 to alanine (T58A) in the endogenous *Myc* locus in the mouse germline. In this way, we generated mice that express MYC protein mutated at T58 but retain the normal genomic regulation of MYC, therefore expression of the *Myc* gene is not de-regulated. Here we report that this mutation causes a tumor-prone phenotype associated with aberrant hematopoietic progenitor cell self-renewal and altered transcriptional regulation of loci proximal to MYC bound enhancers.

## RESULTS

### Generation of mice with a threonine 58 to alanine mutation in endogenous *Myc*

We generated mice with a targeted mutation of Threonine 58 to Alanine (T58A) in the endogenous c-*myc* allele (herein denoted as *Myc*) in mice using standard transgenic techniques. The endogenous *Myc* locus was targeted in ES cells using a construct containing the T58A mutation in exon 2 of *Myc* (Supplemental Fig. S1A). Chimeric mice were generated that transmitted the allele to the germline, and subsequent breeding of founders resulted in a mouse line referred to as *Myc*-T58A. Mice heterozygous for the T58A mutation were used to breed homozygous (*Myc^T58A/T58A^* herein *Myc-T58A),* heterozygous (*Myc^+/T58A^*), and wild-type (*Myc^+/+^*^)^ littermate mice that were used for further experiments. We observed approximately normal mendelian frequencies (Supplemental Fig. S1B), and no evidence of developmental abnormalities in either *Myc^+/T58A^* and *Myc^T58A/T58A^* mice, indicating that mice do not require phosphorylation of MYC at threonine 58 for development.

### *Myc*T58A mouse tissues do not exhibit increased cell cycling or hyperplasia

In the context of ectopic MYC expression, mutation of T58 has been demonstrated to interfere with Fbw7 mediated degradation and augment MYC protein half-life (Welcker et al. 2004; Yada et al. 2004). We measured MYC protein levels by western blot in spleen and thymus to determine the extent to which T58A mutation in the endogenous *Myc* gene alters steady state MYC levels. The level of MYC protein in the *Myc*^T58A/T58A^ knock-in mice was generally elevated approximately 1.5-2-fold in hematopoietic tissues such as spleen and thymus (Fig. 1A). This correlated with an half-life of T58A mutant MYC protein over wild-type MYC from approximately 18 to 30 minutes (Fig. 1A). Increased *Myc* expression, resulting in higher steady-state levels of MYC protein (Fig. 1A), is typically associated with profound increases in cell proliferation. We tested whether the relatively modest increase of MYC levels in c-MycT58A mice might result in increased cell proliferation in different tissues. We observed no evidence of increased proliferation or hyperplasia in lung, brain, colon, small intestine, kidneys, or any hematopoietic organs (Supplemental Fig. S1C,D). Therefore, our MYC-T58A mouse model is distinct from previous mouse models in which increased proliferation and hyperplasia were observed when MYC was deregulated even with relatively low levels of expression (Murphy et al., 2008). These data indicate that the T58A mutant *Myc* gene maintained in its normally regulated, endogenous setting does not elicit the acute proliferation and hyperplasia normally associated with MYC overexpression and genomic deregulation in mice.

**Figure 1.**
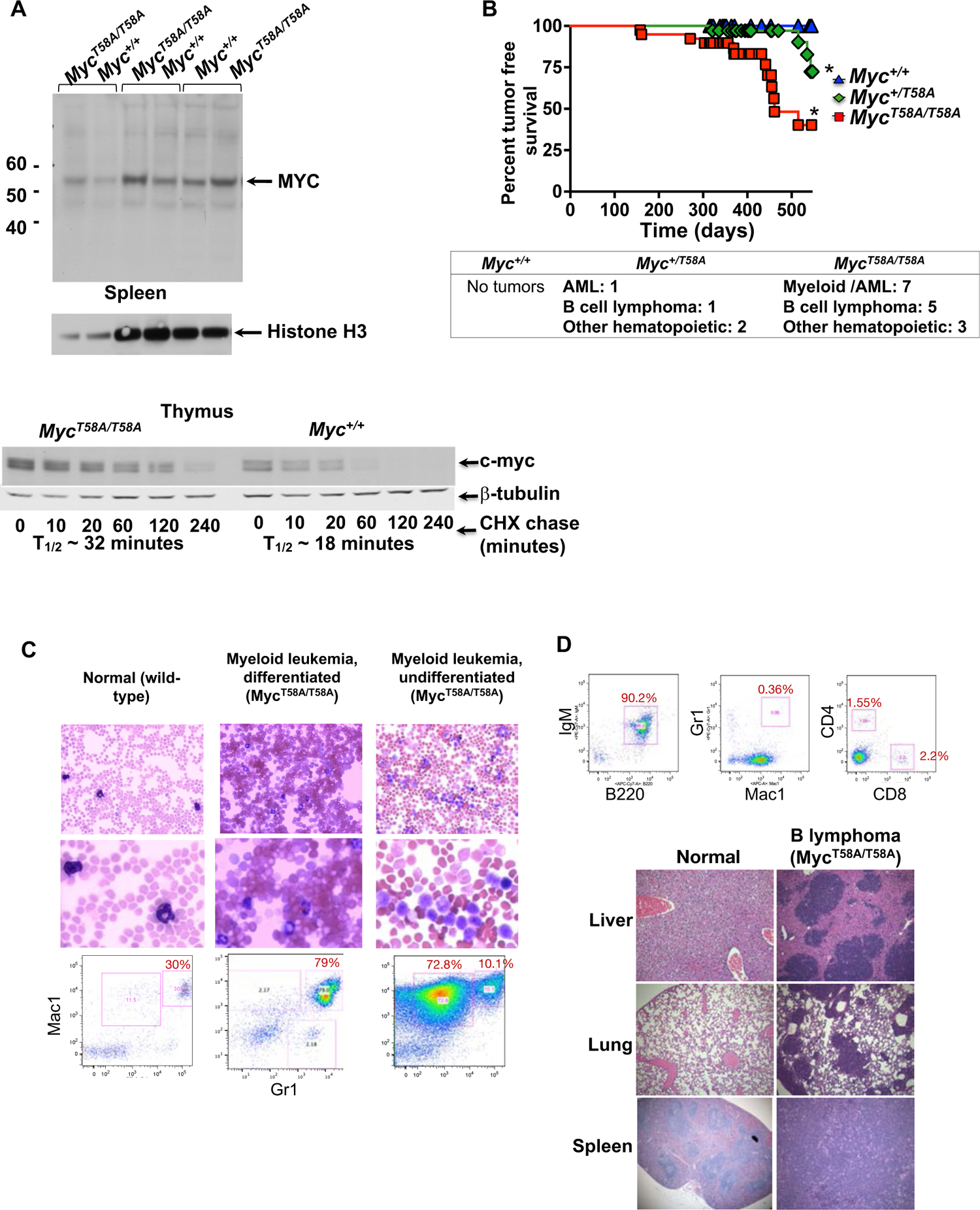
*Myc^T58A/T58A^*mutant mice exhibit increased MYC stability, and are sensitized to late onset hematopoietic malignancies. (A) Protein isolated from whole cell extracts of spleens or thymus from *Myc^T58A/T58A^*and littermate control mice were analyzed by western blotting. Littermate pairs were loaded into adjacent lanes (indicated by brackets), blotted, and blots were detected using antibody to Myc with Histone H3 is used as a loading control (top panel). Single cell suspensions from *Myc^T58A/T58A^* and littermate control (*Myc^+/+^*) mouse thymus were suspended in culture media in the presence of cycloheximide for the time indicated, and western blots for Myc and β-tubulin (loading control) are shown (bottom panel). The calculated half-life (T_1/2_) is indicated. (B) *Myc^T58A/T58A^*and littermate control *Myc^+/+^* mice were allowed to age until the development of disease. Mice were autopsied, and malignancies were confirmed using flow cytometry and pathological analysis. The data shown are a Kaplan-Meier plot of tumor free survival. The asterisk denotes significant p values (p<0.001). A summary of observed malignancies is shown in the table. (C, D) Mice exhibiting a detectable tumor mass or that were moribund were sacrificed, and tissues and blood smears were prepared from normal and malignant mice and stained (Romanowsky). Tissues from malignant animals (liver, lung, and spleen) were stained (Hematoxylin/eosin) for microscopic analysis. Single cell suspensions from malignant *Myc^T58A/T58A^* mice were prepared from blood, bone marrow, and spleen and analyzed by flow cytometry using antibodies specific for the indicated cell surface marker. Cell populations are gated on live cells (DAPI negative).

### Targeting of *Myc* T58A results in sensitivity to late onset hematopoietic malignancies

While we did not observe effects of the MYC-T58A mutation on development or hyperplasia, we nevertheless monitored *Myc^+/+^*, *Myc^+/T58A^*, and *Myc^T58A/T58A^* mice up to 1.5 years of age to determine whether they would develop malignancy with a longer latency. We began to observe some hematopoietic malignancies in *Myc^T58A/T58A^* mice at around 6 months of age, which ultimately affected approximately 60% of these mice by 1.5 years (Fig. 1B). These malignancies were either myeloid or B lymphoid in origin. Myeloid malignancies presented with severe anemia and thrombocytopenia, and malignant cells were characterized by mature or immature myeloid forms expressing myeloid markers (Fig. 1C). Disruption of organ architecture was also seen in marrow, spleen, and often in liver and lung (Supplemental Fig. S1E). Lymphomas were observed in peripheral blood and lymphoid organs (lymph nodes and spleen) and often disseminated into liver and lung (Fig. 1D). These malignant cells expressed B220 and IgM, indicating that they were mature B cell lymphomas (Fig. 1D). Some myeloid and lymphoid malignancies were also seen in the *Myc^+/T58A^* mice, albeit to a lesser extent; indicating that a single, mutant T58A *Myc* allele can sensitize mice to malignancy. Overall, our data indicate that these mice are predisposed to hematopoietic malignancies that require a long latency to develop. Notably this is very different compared with a model in which MYC is overtly over-expressed, such as Eμ-MYC mice in which mice develop lethal tumors by approximately 6 months of age (Eischen et al. 1999).

### Hematopoietic progenitors from *Myc^T58A/T58A^* mice are resistant to apoptosis

The relatively long latency required for tumor formation indicates that malignant evolution in these mice is accompanied by the accumulation of genetic mutations over time. We therefore hypothesized that the mechanism of malignancy in T58A mutant mice is a consequence of some property in their hematopoietic cells that facilitates accumulation of additional genetic events, eventually resulting in the emergence of tumor initiating cells. Importantly, we did not detect increased cell cycling, hyperplasia, or expansion of any particular cell population in any mature hematopoietic lineage, nor did we observe increased cellularity in the marrow, spleen, thymus, or other hematopoietic organs of young or adult *Myc-*T58A mice up to 12 weeks of age (Supplemental Fig. S2A-D).

Hematopoietic precursor cells in the stem or progenitor compartment are typically the cells that become transformed during malignant progression (Krivtsov et al. 2006). Therefore, we tested whether immature hematopoietic stem or progenitor cells from *Myc^T58A/T58A^* mice had any measurable, intrinsic gain of function in biological activity that might be associated with leukemic stem cells prior to the onset of malignancy in young mice (8 weeks old). We observed a modest reduction in the absolute number of phenotypically defined hematopoietic stem cells (HSC’s, Lin-/Sca1+/ckit+/CD150+), indicating a potential consequence of T58A mutation in the stem/progenitor cell compartment (Supplemental Fig. S2E). We therefore assessed whether primitive, reconstituting hematopoietic stem cells (HSC’s) or hematopoietic progenitors from *Myc^T58A/T58A^* mice exhibit increased cell cycling. We found no increase in percent S phase in *Myc^T58A/T58A^* mice as assessed by Ki67 staining using flow cytometry in HSCs, multi-potential progenitors (MPPs), or myeloid lineage restricted progenitors, indicating that these cells cycle at approximately the same rate (Supplemental Fig. S2F).

We next questioned whether T58 phosphorylation of endogenous *Myc* alters the balance between cell survival and apoptosis in hematopoietic precursors. We induced apoptosis in *Myc*T58A-derived cells by culturing cells in the absence of cytokines that are required for hematopoietic cell survival. Bone marrow cells from 8-week-old *Myc*T58A and wild-type control mice were cultured for 24-72 hours in the absence of cytokines to induce apoptosis and were subsequently replated into secondary cultures to measure the loss of hematopoietic progenitor colony forming activity (CFU). We found that, in the absence of cytokines, progenitors from *Myc^T58A/T58A^* mice exhibited approximately two-fold increased survival compared to *Myc+/+* progenitors (Supplemental Fig. S2G). This correlated with a two-fold decrease in apoptosis measured by levels of active caspase 3 in committed myeloid progenitor cells (Supplemental Fig. S2H). We also noticed a modest, but statistically significant, resistance to apoptosis induced by culture in the absence of cytokine in bone marrow derived, pre-B cells from *Myc^T58A/T58A^* mice compared to wild-type controls (Supplemental Fig. S2I,J). Together these data indicate that phosphorylation of MYC T58 contributes to regulation of apoptosis in hematopoietic progenitors during normal hematopoiesis.

### Hematopoietic progenitors in *Myc^T58A/T58A^*mice exhibit aberrant self-renewal activity

The hallmark of HSC activity is the ability to self-renew, a trait that is often acquired by leukemic cells in non-self-renewing progenitor populations (Krivtsov et al., 2006). We hypothesized that the MYC-T58A mutation potentiates self-renewal in hematopoietic progenitors that normally do not possess self-renewal activity, thus providing these cells with aberrant activity that allows them to emerge as leukemic precursors. To address this possibility, we cultured hematopoietic progenitors *in vitro* in clonogenic assays, and assessed the ability of daughter cells from these colonies to self-renew by re-plating them in secondary cultures. Hematopoietic progenitors derived from *Myc^T58A/T58A^* mice formed colonies in secondary culture at a 3-fold higher frequency than progenitors from wild-type littermate control mice (Fig. 2A). These *in vitro* data provide evidence of an abnormal, gain of function in self-renewal of *Myc^T58A/T58A^* hematopoietic progenitors. To measure the self-renewal capacity of hematopoietic stem and progenitor cells more definitively i*n vivo*, we sorted hematopoietic precursor populations from *Myc^T58A/T58A^* and wild-type littermate control mice and compared the ability of distinct stem and progenitor cells to reconstitute lethally irradiated recipients (Fig. 2B). Sorted hematopoietic stem (HSCs), multi-potential progenitors (MPPs) and committed myeloid progenitors (CMPs) were transplanted at dilution along with “competitor” cells (expressing distinct CD45 antigens) and analyzed over time in recipient mice for the presence of cells derived from the transplanted progenitors. HSCs are unique in their capacity to reconstitute the hematopoietic system of lethally irradiated recipient mice long-term, while multi-potential progenitors and committed myeloid progenitors do not possess this ability. Strikingly however, *Myc^T58A/T58A^*MPPs reconstituted lethally irradiated mice up to at least 19 weeks post-transplant, indicating aberrant long-term self-renewal activity (Fig. 2C,D). Importantly we observed both myeloid and lymphoid long-term reconstitution in these mice. As expected, MPPs derived from wild-type control littermate mice exhibited no self-renewal. Taken together, the evidence indicates a striking aberrant self-renewal of MYC-T58A-mutant hematopoietic progenitors that is likely to be a key factor in sensitizing these mice to myeloid neoplasia. Furthermore, both the *in vitro* culture and *in vivo* transplant data indicate that this phenotype is intrinsic to hematopoietic progenitors, and not a consequence of changes in the bone marrow niche or microenvironment.

**Figure 2.**
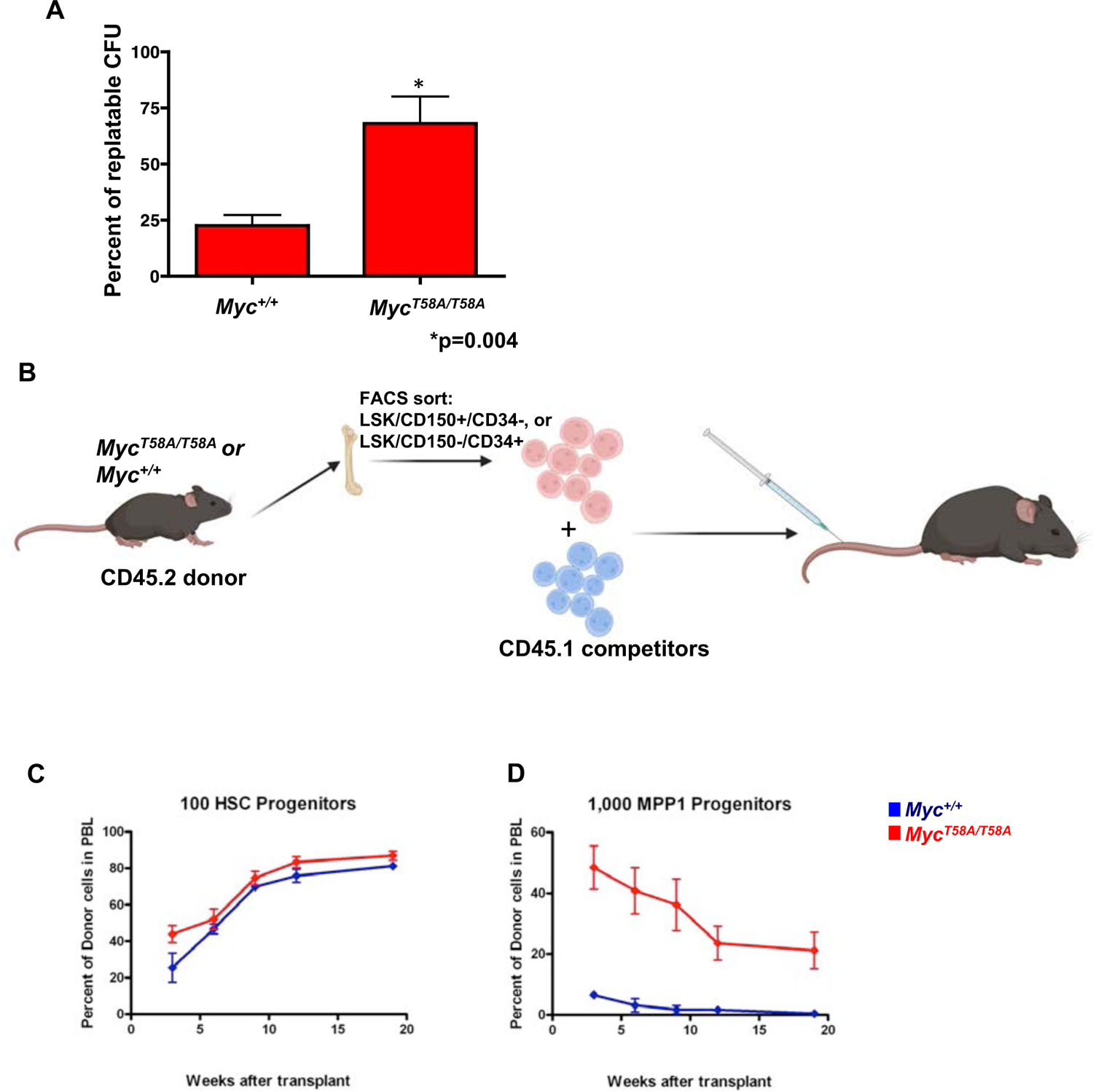
Aberrant self-renewal of hematopoietic progenitors from *Myc^T58A/T58A^* mice. (A) Bone marrow hematopoietic progenitors were cultured from either *Myc^T58A/T58A^* mice or *Myc^+/+^* mice in clonogenic methylcellulose assays in the presence of cytokines to support the growth of myeloid progenitor colonies. After 10 days in culture, individual colonies were picked and replated as secondary cultures supplemented with cytokines. Secondary colony formation was scored at day 10 as replatable CFU, and the ratios of replatable colonies compared to total colonies plated are shown (y axis). The data shown are representative of two independent experiments. (B, C) Hematopoietic stem and progenitor cells from the bone marrow of 3 pooled *Myc^T58A/T58A^*mice and *Myc^+/+^* control littermates were sorted by flow cytometry. Hematopoietic stem cells (HSC), and one population of multi-potential progenitors (called MPP1) were sorted based on expression of the indicated cell surface markers. The sorted stem or progenitor cells (CD45.2) were then mixed at decreasing cell doses with competitor cells (CD45.1) and transplanted into lethally irradiated recipient mice. Antibodies specific to the two CD45 isoforms were used to determine the levels of reconstitution from the indicated sorted cell populations. The data are shown as percent CD45.2 donor.

### Identification and characterization of hematopoietic progenitors with increased self-renewal in MYC-T58A mutant mice

To further identify and characterize hematopoietic progenitor populations in MYC-T58A mutant mice, we used single cell methods to profile the transcriptional outcome and epigenetic state of hematopoietic stem and progenitor cells in 8-week-old mice, prior to the onset of any disease. Lin^-^Sca1^+^ckit^+^ cells from littermate WT and *Myc^T58A/T58A^* mice were obtained and sorted from bone marrow in two replicate experiments. These cells were processed using single cell capture and barcoding for multiome RNA-seq and ATAC-seq (Supplemental Fig. S3A).

The results for captured WT and T58A single cells were pooled in the subsequent analysis and the datasets were filtered for high quality cells and integrated to account for technical variability (Butler et al. 2018; Hafemeister and Satija 2019; Stuart et al. 2021). Dimension reduction and clustering of these cells revealed the existence of 14 components, comprising major stem and progenitor subsets. Most of these subsets were easily identified based on the expression of defining genes for different lineages (see METHODS, and Supplemental Fig. S3B). We further refined these progenitor subsets based on recently described analytical approaches and single cell data (Cabezas-Wallscheid et al. 2014; Eaves 2015; Nestorowa et al. 2016; Belluschi et al. 2018; Rodriguez-Fraticelli et al. 2018; Konturek-Ciesla et al. 2023). There were no marked changes in the relative number or percent of any defined progenitor type upon integration of the WT and T58A data (Fig. 3A, Supplemental Fig. S3C). Lymphoid and myeloid populations ordered in pseudo-time indicated that T58A progenitors did not have an obviously compromised differentiation status (Fig. 3A).

**Figure 3.**
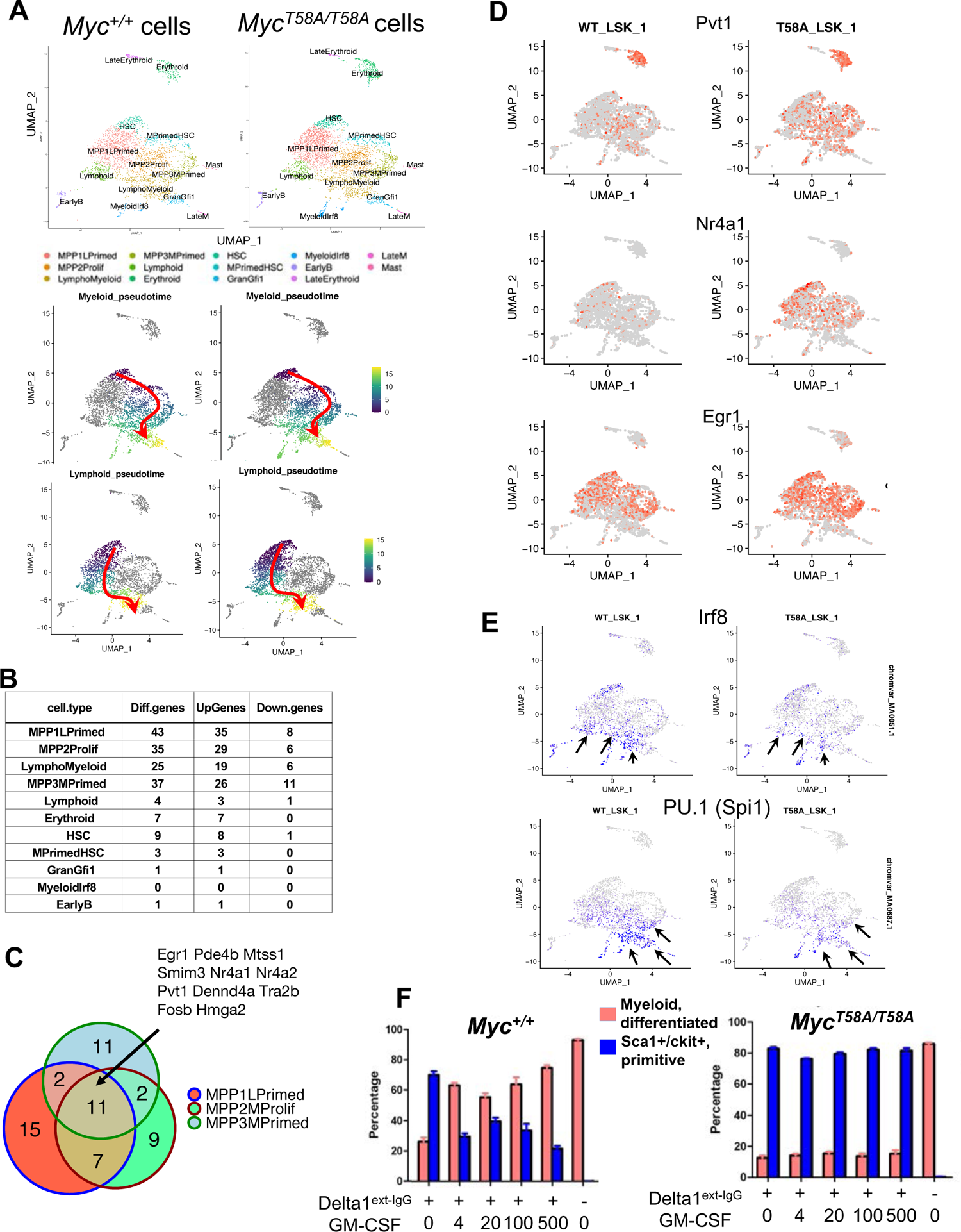
Single cell multiome analysis comparing *Myc^+/+^* and *Myc^T58A/T58A^*hematopoietic stem and progenitor cells. (A) Hematopoietic stem and progenitors (LSK sorted cells) from freshly isolated bone marrow of either *Myc^+/+^*or *Myc^T58A/T58A^* mutant Myc mice were sorted and barcoded to identify single cells using the 10X Genomics platform for single cell multiome (RNA-Seq and ATAC-Seq) analyses. Two mice per genotype were used in two independent experiments to generate 4 datasets (4 mice per genotype, for a total of 8 mice). After alignment and analysis using R packages Seurat and Signac (based RNA content and removal of duplicate barcodes), 7929 cells were included in the analysis. The datasets were then integrated (to account for batch effects), and clustered based on single cell RNA-Seq of *Myc^+/+^* and *Myc^T58A/T58A^*. Following dimension reduction, progenitor cell populations were identified and labeled as detailed in METHODS. Primitive hematopoietic progenitor cells were ordered in pseudo time using the R package Monocle3. HSC’s were set at 0, and cells were ordered along either lymphoid or myeloid pseudotimes. (B) The number of differentially expressed (Diff.genes), Up-regulated (Up.Genes), and Down-regulated (Down.Genes) comparing T58A mutant to WT cells, determined for each population of progenitor cells. Differentially expressed genes were determined by Wilcoxon rank sum analysis (p < 0.05). (C) Venn diagram depicting differentially expressed genes in 3 different hematopoietic progenitor cell populations (MPP1_LPrimed, MPP2_Prolif, and MPP3_MPrimed). A set of 11 Myc target genes are shown that were up-regulated in *Myc^T58A/T58A^* mutant cells in all analyzed progenitor cell populations. (D) Heatmap of single cells showing the expression of selected genes found to be up-regulated in *Myc^T58A/T58A^* mutant progenitor populations. (E) Consensus binding motif activity in single cell ATAC-Seq in progenitors in *Myc^+/+^* and *Myc^T58A/T58A^*mutant mice. (F) Hematopoietic progenitor cells were purified from bone marrow of *Myc^+/+^* or *Myc^T58A/T58A^*mice (LSK sorting). Sorted cells were then cultured in wells coated with Delta1 ligand to activate the Notch pathway in the presence of cytokine cocktail (see METHODS). After two weeks of culture, increasing amounts of GMCSF was added to the cultures to promote myeloid differentiation. Differentiation was assessed by flow cytometry staining for Sca1/ckit (Sca1/ckit+, primitive) which is retained on primitive cells, and Mac1/Gr1/F480 (Myeloid, differentiated) to identify differentiated cells.

We next asked whether the multi-potential progenitor subsets in T58A mice with aberrant self-renewal (Fig. 2C) exhibit a differential gene expression signature. We compared gene expression in WT vs T58A progenitor cell populations in the integrated data. This analysis revealed that 139 genes were differentially expressed over all T58A mutant stem and progenitor cell populations (by Wilcoxon Rank Sum test, adjusted p value < 0.05). The fold changes in gene expression were relatively modest, and the overwhelming majority of differentially expressed genes were observed in the primitive and multi-potential progenitors, with fewer changes in the most primitive HSC compartment or relatively mature progenitors (Fig. 3B,C). Differentially expressed genes were enriched for pathways that regulate hematopoiesis, particularly inflammation (such as TNF, NFκB signaling), and differentiation (Foxo1 and Spi1 target genes) (Supplemental Fig. S3D). We found increased expression of important transcription factors that mediate HSC and leukemic cell self-renewal, including nuclear hormone receptors (Nr4a1, Nr4a2), Egr family members (Egr1, Egr3), and chromatin modifiers (Kdm6b, Hmga2); many of which are known to be regulated by MYC (Fig. 3C, D, Supplemental Fig. S3E) (Scheicher et al. 2015; Mallaney et al. 2019; Desterke et al. 2021; Sheng et al. 2021). We also noted increased expression of Pvt1, which is long, non-coding RNA often co-amplified with MYC in oncogenesis and associated with increased MYC stability and expression (Tseng et al. 2014; Cho et al. 2018). Though the expression of these genes varied depending on the population, the changes were particularly evident in the most proliferative progenitor compartments (MPP-L-primed, MPP-M-primed, and MPP-prolif), with a core set of 11 genes up-regulated in all 3 of these populations (Fig. 3C). This is consistent with the observation that Myc expression is highest in these cell populations (Supplement Fig. S3B). Overall, these data indicate that multi-potential progenitors (with aberrant self-renewal capacity) in the marrow of *Myc^T58A/T58A^* mice have only mild, if any, disruptions in differentiation status prior to onset of detectable disease. They do however exhibit consistent differential expression of MYC-regulated genes known to be important in differentiation, inflammation, and maintenance of the primitive hematopoietic compartment.

The chromatin accessibility of each progenitor population, as identified based on RNA-seq clustering, was assessed using the associated ATAC-seq data. To determine whether accessible regions had altered transcriptional activity, we performed DNA binding motif analysis using Chromvar to assign a motif score to every cell for all motifs in the JASPAR database (Schep et al. 2017) based on accessible peaks. Interestingly, the MYC consensus binding motif (CACGTG) single cell score was not increased in MYC-T58A cells relative to wild-type cells (Supplemental Fig. S3F). We did however note alterations in motifs important in inflammation and differentiation, such as IRF8 and PU.1. These motif scores were diminished in T58A mutant cells, as was evident in relatively differentiated myeloid and lymphoid progenitors (Fig. 3E). This indicates that T58A progenitors may be less poised for both myeloid differentiation and the induction of inflammatory processes during differentiation.

Based on these results, we reasoned that T58A mutant progenitor cells could possess a decreased propensity for differentiating into inflammatory myeloid cells. Indeed, when LSK sorted, primitive hematopoietic stem and progenitor cells were cultured in delta ligand to activate the Notch pathway (which promotes primitive lymphopoiesis), T58A mutant cells ultimately accumulated larger numbers of primitive hematopoietic cells expressing ckit and Scal (Supplemental Fig. S3G) after 2 weeks of culture compared to wild-type cells (5×10^9^ total T58A cells compared to 1×10^8^ WT cells). When cells were cultured with delta ligand in the presence of increasing amounts of GM-CSF to promote myeloid differentiation, wild-type cells predominantly differentiated, as over 75% of cells began to express myeloid markers, and less than 20% retaining Sca1 and ckit expression at the highest concentration of GM-CSF (Fig. 3F). Strikingly, only about 15% of T58A cells differentiated into myeloid cells, while over 80% retained Sca1/ckit expression even at the highest concentration of GM-CSF (Fig. 3F), although they differentiated normally in the absence of Delta ligand.

Overall, these data characterize a defined set of progenitors in T58A mice that have promiscuous self-renewal capacity prior to malignant progression. Alterations in the self-renewal signature is evident particularly in multi-potential progenitors from these mice, consistent with the aberrant self-renewal observed in transplant and re-plating studies (Fig. 2). Moreover, we find that the T58A progenitors are functionally distinct relative to wildtype progenitors in that, while not present as an overtly expanded population in steady state hematopoiesis, they nonetheless appear to have a diminished capacity for myeloid differentiation and inflammation when stimulated in culture.

### B lymphocytes from *Myc^T58A/T58A^* mice have an altered transcriptome relative to WT lymphocytes

While a subset of *Myc^T58A/T58A^* mice developed myeloid leukemia, a significant number of *Myc^T58A/T58A^*mice developed B cell lymphomas (5/12 neoplasms) characterized by B220 and IgM expression (Fig. 1B, D). B lymphoid progenitors (Pre-B) cells from the bone marrow of *Myc^T58A/T58A^*mice were resistant to apoptosis and responded differentially to IL7 stimulation, (Supplemental Fig. S2I,J; Supplemental Fig. S4A). To better understand the events leading to lymphomagenesis in these mice we asked what transcriptional alterations are manifested in the B cell lineage. To this end, we assessed the transcriptional profile of bulk populations of B cells derived from littermate *Myc+/+* and *Myc^T58A/T58A^* mice by comparing B cells derived from bone marrow and spleen using RNA-Seq following mitogen stimulation for 48 hours.

In IL7 stimulated Pre-B cells we identified 520 genes with increased expression and 557 with decreased expression, comparing T58A to WT controls, (adjusted p value < 0.05, Fig. 4A, Supplemental Fig. S4B). Genes with increased expression in *Myc^T58A/T58A^*mice were significantly enriched for known MYC target genes (Fig. 4B). Many of these up-regulated genes were associated with metabolic pathways known to be involved in MYC activation, including glucose metabolism, the mTOR pathway, and nucleotide and fatty acid biosynthesis (Fig. 4B). Importantly, other metabolic pathways activated by MYC in B cells (such as oxidative phosphorylation) were not found to be increased in T58A mutant cells. Hence, cells expressing T58A mutant MYC selectively increased the expression of a subset of MYC target genes.

**Figure 4.**
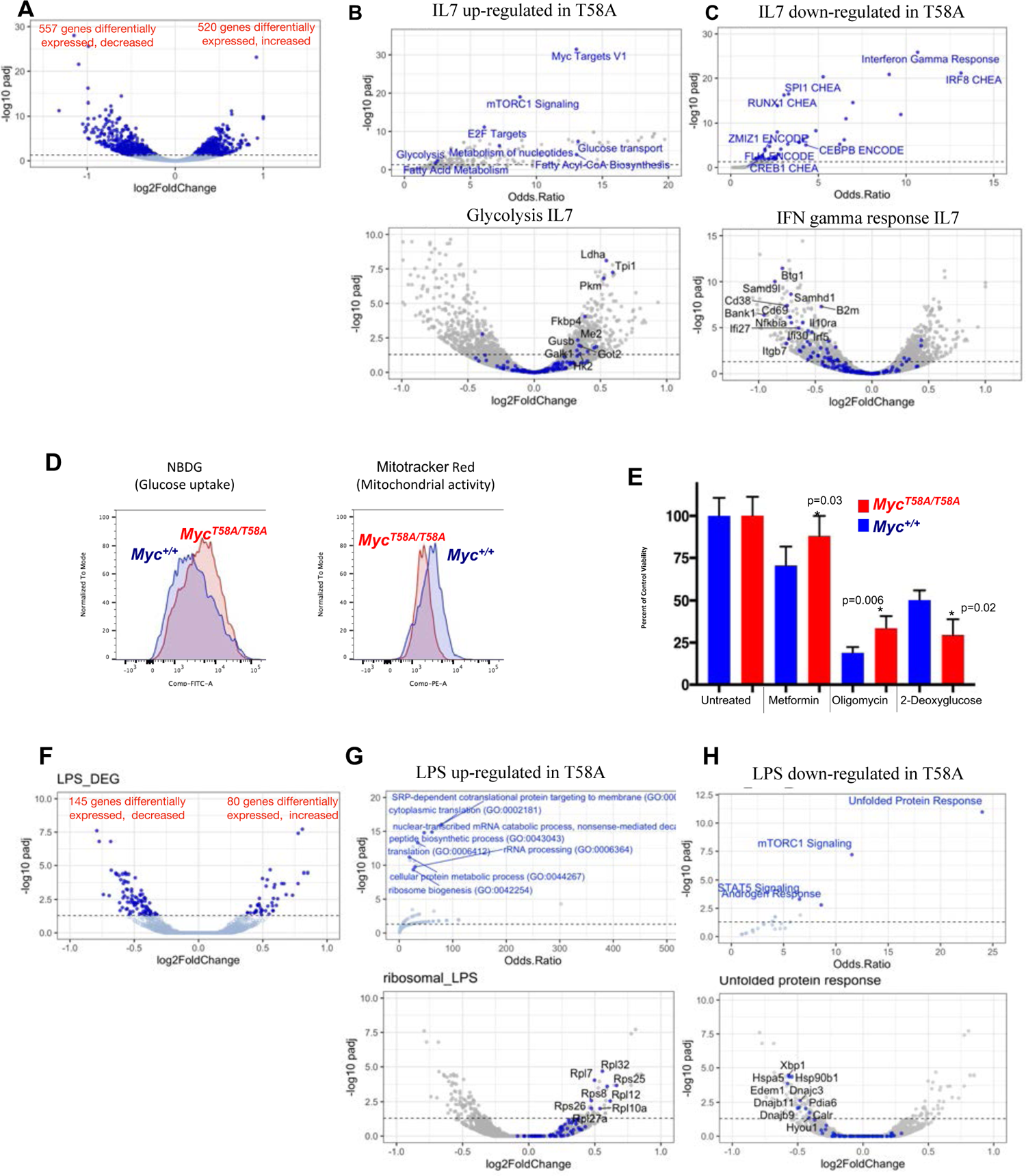
RNA-Seq analysis comparing *Myc^+/+^* to *Myc^T58A/T58A^* mutant Pre-B and mature B cells. (A-C) B cells were sorted from paired, littermate WT or T58A mutant mouse bone marrow. RNA was extracted from cells after stimulation with IL-7 for 48 hours, and libraries were prepared and barcoded. After sequencing, analysis was performed using the DESeq2 R package. Differentially expressed genes are depicted in dark blue on the volcano plot in panel A, with each gene plotted as the log2 fold-change (x-axis) against the -log10 transformation of the p value (y-axis). Enrichment analysis (using EnrichR package) was performed on up regulated (panel B) and down regulated (panel C) genes. The panel B plot (top) depicts enriched pathways from MSigDB Hallmark and Reactome genesets, and (bottom) a volcano plot showing differentially expressed genes involved in glycolysis labeled and highlighted in blue. Panel C (top) shows enrichment of down regulated genes for pathways (MSigDB Hallmark and ENCODE/CHEA ChIP bound transcription factors), and (bottom) differentially expressed genes involved in interferon response, which are labeled and highlighted in blue. (D) Flow cytometry analysis of glucose uptake potential was assessed by incorporation into cells of fluorescent labeled NBDG (right panel) and Mitotracker red (left panel) in *Myc^+/+^* and *Myc^T58A/T58A^*mutant cells stimulated with IL-7. The fluorescent intensity (x-axis) is plotted against cell number (y-axis) in the plot overlays, with the curve for *Myc^+/+^* cells in blue, and *Myc^T58A/T58A^* mutant in red. (E) IL-7 stimulated cells were untreated, or treated with either mitochondrial inhibitors (Metformin, Oligomycin) or glucose uptake inhibitor, 2-DG (x-axis). Cells were counted, and plotted as percent of control, untreated cells (y-axis). *Myc^+/+^* cells are shown in blue, and *Myc^T58A/T58A^* cells in red. (F-H) B cells were sorted from *Myc^+/+^* or *Myc^T58A/T58A^* mutant mouse spleens using anti-B220 coated magnetic beads (MACS). RNA was extracted after stimulation with LPS for 48 hours as described above. Paired analysis was performed using the DESeq2 R package. Up and down regulated genes are shown in dark blue on the volcano plot in panel F, and each gene is plotted as log2 fold-change (x-axis) against the -log10 transformation of the p value (y-axis). Enrichment analysis (EnrichR) was performed for up regulated (panel G) and down regulated (panel H) genes. The panel G plot (top) depicts enriched pathways (from GO biological process), and (bottom) a volcano plot showing differentially expressed genes involved in ribosomal protein biogenesis labeled and highlighted in blue. Panel H (top) shows enrichment of down regulated (MSigDB Hallmark genesets), and (bottom) differentially expressed genes involved in the unfolded protein response, which are labeled and highlighted in blue.

Differentially expressed genes with relatively decreased expression upon T58A mutation were enriched for inflammatory pathways, especially those associated with interferons and activation by IRF8, NFκB, and Spi1/PU.1 (Fig. 4C, Supplemental Fig. S4C). We also observed down-regulation of MHC class II genes (Supplemental Fig. S4B). Moreover, apoptosis-associated genes were also decreased in T58A mutant Pre-B cells (Supplemental Fig. S4C), consistent with our findings that these cells are resistant to apoptosis (Supplemental Fig. S2I,J). In particular, transcription of the pro-apoptotic effector Bcl2l11 (BIM), a well-established MYC target gene (Muthalagu et al. 2014) that is activated in B cells, was found to be decreased in T58A cells (Supplemental Fig. S4D); in agreement with previous studies of this mutation in the context of MYC over-expression (Hemann et al. 2005).

The RNA-Seq results suggested that T58A mutant cells are metabolically altered to be more dependent on glucose. In agreement with this, we found that, relative to wild-type cells, T58A mutant Pre-B cells exhibited increased uptake of the glucose analog 2-NBDG, indicating a propensity for increased glucose flux, accompanied by an evident decrease in mitochondrial activity (Fig. 4D). Notably, expression of genes encoding rate-limiting enzymes important for glucose metabolism (Ldha, Hk2) were increased in T58A cells (Fig. 4B, Supplemental Fig. S4D). Consistent with these observations, while T58A cells were relatively resistant to the effects of mitochondrial complex I and V inhibitors (metformin and oligomycin, respectively), they were hypersensitive to blockage of glucose uptake (2-deoxyglucose) (Fig. 4E). Together these data indicate that T58A Pre-B cells have an increased rate of glycolysis uncoupled from augmented mitochondrial oxidative phosphorylation, suggesting utilization of the glucose-derived carbon equivalents for increased biosynthesis (nucleotide, fatty acid metabolism). Decreased activation of inflammatory and apoptotic pathways are also consistent with the increased survival and self-renewal of T58A mutant cells.

The transcriptional response of mature B cells to LPS stimulation is profoundly MYC dependent, as MYC is known to be involved in the activation of thousands of genes, many of which are hallmarks of MYC expressing cells (Nie et al. 2012; Kieffer-Kwon et al. 2017; Tesi et al. 2019). Because T58A mutant cells survive and proliferate at a higher rate relative to wild-type cells in LPS culture (Supplemental Fig. S4E), we asked whether we might detect relevant transcriptional changes in this model. We identified a very specific signature from a relatively small number of genes (80 genes up-regulated, and 145 genes down-regulated) that were differentially expressed in mature B cells from T58A vs. WT mice stimulated with LPS (Fig. 4F, Supplemental Fig. S4F). Genes involved in regulation of protein translation were highly enriched among the set of up-regulated genes, and ribosomal proteins comprised a significantly increased component of this response (Fig. 4G). Strikingly, the major class of down-regulated genes in LPS stimulated B cells were those involved in the unfolded protein response (UPR) (Fig. 4H). These data further indicate that T58A mutation increases the expression of only a small fraction of MYC activated genes after LPS stimulation. Moreover, as the UPR is known to be a MYC dependent process (Babcock et al. 2013), we surmise that the T58A mutation causes the transcription of these genes to be attenuated compared to WT MYC.

Taken together, the data indicate that T58A mutant MYC alters gene expression of a limited subset of MYC target genes in primitive and mature B cells. These changes in gene expression are quite consistent with the increased survival and metabolic re-wiring of these cells, and some MYC target gene RNAs were predictably increased. However, many MYC target genes involved in inflammation, apoptosis, and unfolded protein response were decreased in T58A mutant cells.

### Analysis of MYC occupancy, RNA polymerase II distribution and enhancer proximity at differentially expressed genes

Our results show that only a subset of canonical MYC target genes expressed in B lymphocytes are increased or decreased in cells with the T58A mutation, indicating that this mutation primarily affects a subpopulation of genes within the MYC transcriptional program. We anticipated that increased amounts of MYC would occupy promoters of up-regulated genes, since MYC generally activates transcription. To test this, we performed genomic analyses in Pre-B cells (stimulated with IL7) and mature B cells (stimulated using LPS) to identify genomic loci occupied by MYC in WT compared to T58A cells. As expected, we found widespread MYC binding at gene promoters in WT cells, particularly in expressed genes, and in many differentially expressed genes (Fig. 5A,B). Increased occupation of promoters by T58A mutant MYC was detected in both LPS and IL-7 stimulated B cells, consistent with increased levels of MYC in these cells (Fig. 5C). However, MYC genomic occupation was generally increased regardless of whether genes exhibited increased, decreased, or unchanged expression levels in MYC-T58A mutant cells. Hence, the gene expression changes are not explained solely by increased amounts of MYC-T58A on promoters. Moreover, clustering of Myc coverage at promoters indicated that the MYC-T58A mutant does not occupy promoters in addition to those bound by wild-type MYC, rather the increased MYC occupancy occurs at the same loci bound by wild-type MYC (Supplemental Fig. S5A).

**Figure 5.**
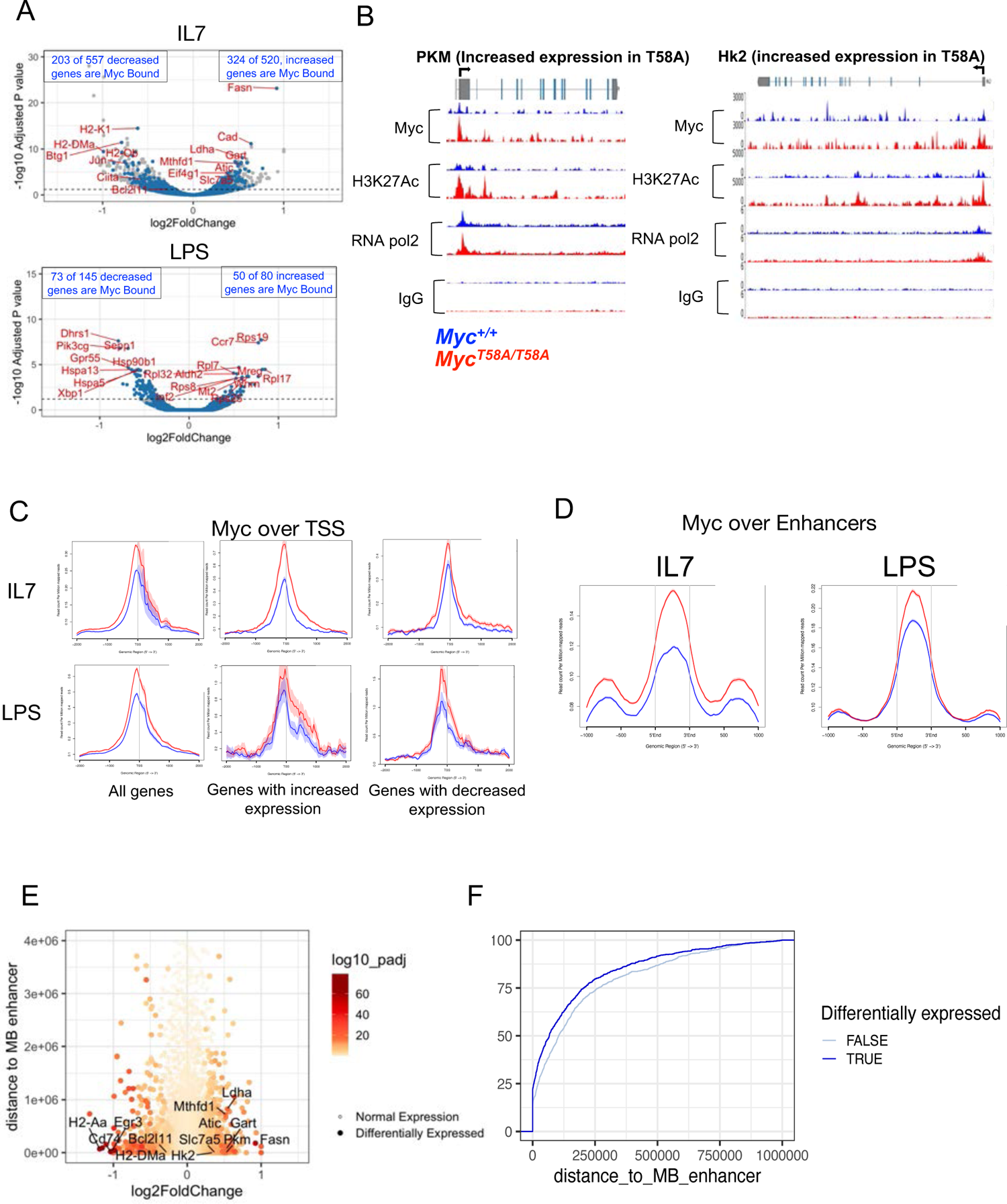
Genomic analysis of Myc at promoters and enhancers in *Myc^+/+^* vs *Myc^T58A/T58A^*cells. Pre-B or mature B cells sorted from bone marrow and spleen of *Myc^+/+^*and *Myc^T58A/T58A^* mice were stimulated with IL-7 (for marrow derived Pre-B cells, upper panels) and LPS (mature B cells, lower panels) for 48 hours, and 1 million cells were used for Auto Cut&Run to detect Myc, H3K27Ac, and H3K3Me2, followed by barcoding, sequencing and alignment. (A) MYC bound genes are shown (dark blue circles) on the volcano plots depicting the RNASeq datasets (also shown in Figure 4) comparing gene expression in *Myc^T58A/T58A^*vs *Myc^+/+^* B cells stimulated with IL7 (left panel) and LPS (right panel). Each gene is plotted as the log2 fold-change (x-axis) against the -log10 transformation of the p value (y-axis). Differentially expressed genes scatter above the dotted horizontal line (padj > 0.05). Genes labeled in red are a sample of differentially expressed genes. (B) Genomic tracks showing MYC, H3K27Ac, Pol2, and control, IgG peaks enriched in Cut&Run and ChIP-Seq (for Pol2) for *Hk2* and *Pkm* genomic loci which have increased expression in *Myc^T58A/T58A^* mutant cells. Tracks for *Myc^+/+^* cells are shown in blue, and *Myc^T58A/T58A^*are shown in red. (C) Plots of normalized read counts (y-axis) were generated centered on the TSS of all genes (left panel), genes with increased expression (middle panel), and genes with decreased expression (right panel). Plots are shown for WT cells (blue), and T58A cells (red). (D) Enhancers were identified in WT and T58A mutant, IL-7 stimulated Pre-B (IL-7) and LPS stimulated mature B cells by peakcalls for H3K27Ac and H3K4Me2 Cut&Run (see Supplemental Fig S5F). Enhancers were defined as sites enriched for both marks (and not promoter bound). Myc Cut & Run signal was plotted over all identified enhancer peaks for *Myc^+/+^* (blue lines) and *Myc^T58A/T58A^* (red lines) cells. (E) Enhancers that were Myc bound were identified by Myc peak calls that overlap enhancers. The distance to the nearest Myc bound enhancer for every gene was calculated. The log2FoldChange for every gene (determined by RNA-Seq comparing WT to T58A in IL-7 stimulated Pre-B cells) was then plotted (x-axis) against the distance to the nearest Myc bound enhancer (y-axis). Differentially expressed genes are represented by filled circles, and the adjusted p values determined for each gene by RNA-Seq is shown by red color intensity. (F) Cumulative distribution function comparing the distance of each gene to the nearest Myc bound enhancer. Genes were ranked according to distance to the nearest Myc bound enhancer. Differentially expressed (dark blue line) are compared to an equal sized set of random genes that are expressed during IL7 treatment, but not differential (light blue line). Statistical correlation assessing the difference between the curves was determined by Kolmogorov-Smirnoff test (p = 8×10^-6^).

Because overexpressed MYC has been shown to promote both transcription initiation and elongation during gene expression we performed ChIP to measure the distribution of RNA polymerase II (Pol2). Genes exhibiting increased expression in T58A cells contained more overall Pol2 reads at both promoters and gene bodies, consistent with their elevated expression (Supplemental Fig. S5B). As MYC transcriptional function has been shown to be associated with control of Pol2 pause/release, we determined the Pol2 traveling ratio by comparing promoter-occupied to gene body-occupied RNA Pol2 (Supplemental Fig. S5 B-D). Surprisingly, traveling ratios were not significantly altered in MYC-T58A compared to WT MYC cells, indicating that the MYC T58A mutation does not overtly affect the ratio of promoter-bound to elongating transcripts. Hence, the increased expression of genes seen for T58A mutant Myc may be explained by modestly increased RNA Pol2 loading on promoters and subsequent Pol2 precession through gene bodies in addition to increased H3K27 acetylation (Supplemental Fig.S5 B-D,F; Fig 5B).

We noted that differentially expressed genes were often clustered on the genome (e.g., Supplemental Fig. S5E) prompting us to consider the possibility that differential gene expression due to T58A might be related to the proximity of the clusters to MYC-bound enhancers. We therefore employed CUT&RUN to identify enhancers in our WT and T58A mutant cells. Our criteria for enhancer identification included non-promoter associated peaks containing both H3K27Ac and H3K4Me2 histone marks and MYC peaks (Supplemental Fig. S5F). Similar to promoters, we found elevated levels of MYC T58A occupation at enhancers (Fig. 5D), again consistent with increased MYC levels in these cells. Interestingly, gene expression changes (comparing WT and T58A mutant cells) were highly dependent on proximity to MYC-bound enhancers in that genes that reside closer to mapped MYC-bound enhancers were significantly more likely to be differentially expressed (Fig. 5 E,F; Supplemental Fig. S5G). Furthermore, this association was evident regardless of whether differentially expressed genes were occupied by Myc (Supplemental Fig. S5H). We collected all the Myc-bound enhancers that were in close proximity (100 kb) to at least one differentially expressed gene in IL7 stimulated Myc-T58A cells. We found that 444 Myc bound enhancers were proximal to differentially expressed genes, of which 50 enhancers were proximal to >2 differentially expressed genes (Supplemental Table S1). Correspondingly, approximately half of the differentially expressed genes were proximal (within 100 kb) to Myc-bound enhancers. Myc-bound enhancers were often found in introns of differentially expressed genes (Supplemental Fig. S5G), such as in the Bim (Bcl2l11) locus. H3K27Ac was also increased at MYC-bound enhancers in Myc-T58A mutant cells, consistent with increased active chromatin (Supplemental Fig. S5F). We surmise that the increased, and potentially prolonged, association of MYC T58A, compared to WT MYC, with both enhancers and enhancer-proximal genes results in altered transcription, and increased loading of RNA Pol2 on genes with increased expression (Fig. 6).

**Figure 6.**
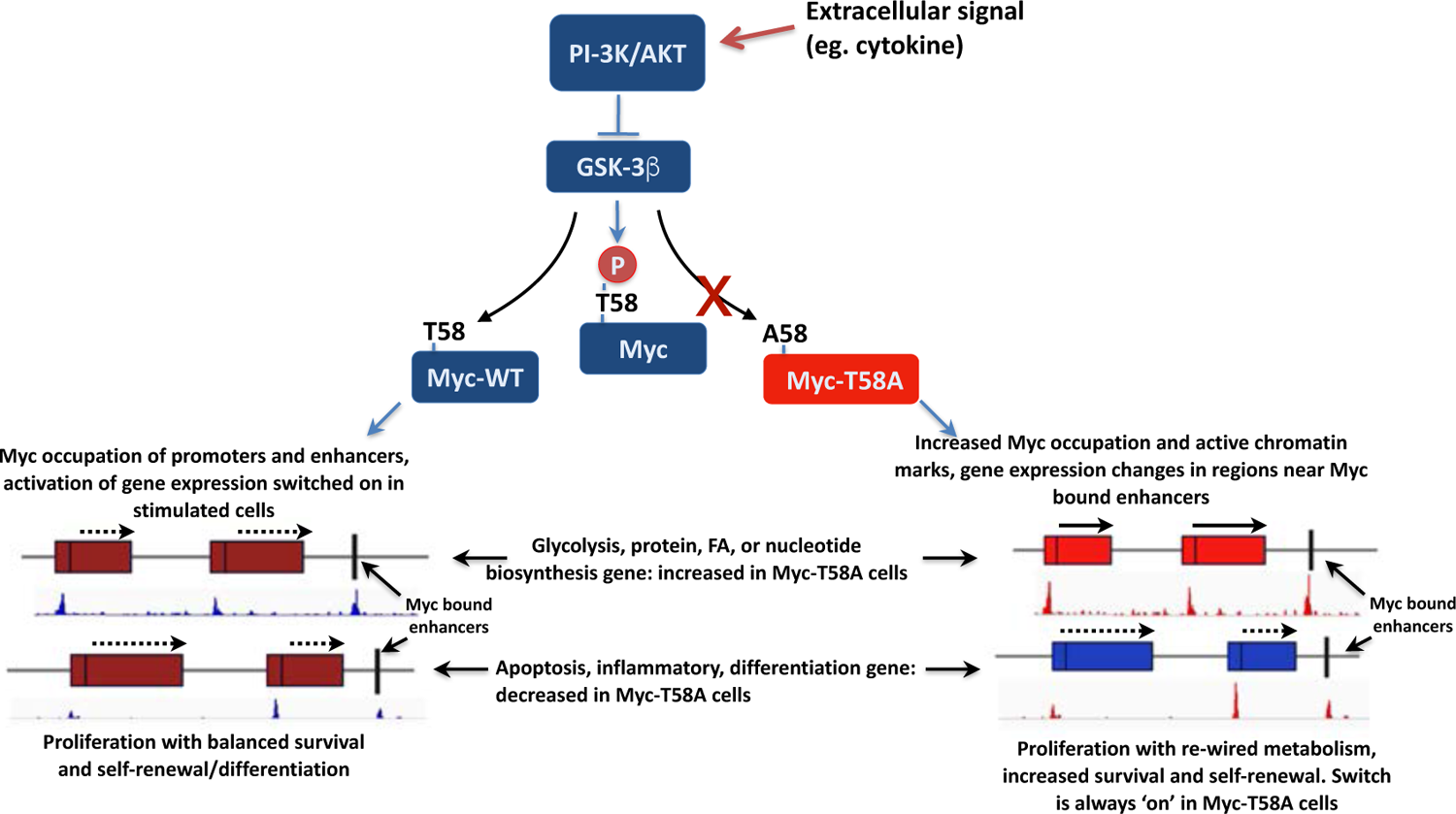
Model depicting dynamics of transcription as a consequence of Myc-T58 phosphorylation. MYC is activated by extracellular signaling via PI-3K and AKT activation, which inhibit GSK3β, and Myc-T58A phosphorylation. This activates a molecular ‘switch’ that changes the state of cells, allowing increased proliferation with balanced survival and self-renewal. With T58A mutation, the switch is locked into the on position changing transcription of regions near MYC bound enhancers. Many genes involved in glycolysis, nucleotide and fatty acid biosynthesis, and protein translation genes (shown in red) are increased in expression in T58A mutant cells, while many inflammatory, apoptosis, and differentiation genes are decreased (shown in blue).

## DISCUSSION

We have developed and characterized a mouse model in which a regulatory, lymphoma associated phosphorylation site in c-MYC was inactivated by mutating its normally expressed, genomic allele. This single amino acid mutation is sufficient to predispose mice to leukemia and lymphoma with a relatively long latency in the absence of genetic deregulation or high levels of MYC over-expression. Pre-malignant hematopoietic progenitors and B cell precursors harboring this mutation had enhanced self-renewal and resistance to apoptosis associated with transcriptional perturbations of MYC target genes. Interestingly, the effects of this point mutation appear confined to the hematopoietic system, highlighting the importance of this phosphorylation in regulating normal hematopoiesis. This is consistent with the essential role of MYC in normal hematopoiesis (Laurenti et al. 2008) as well as the association and severe biological outcome of MYC Box I mutation in B cell lymphomas, and the more recent observation of this mutation in myeloid leukemias (Ferraro et al. 2021). When MYC is overexpressed or overtly de-regulated in mice, affected tissues undergo an initial phase of hyperplasia prior to tumor onset. This is accompanied by increased levels of apoptosis, increased cell cycling and changes in metabolism. Notably, we did not find any evidence of hyperplasia, increased cell cycling, or apoptosis in the c-MycT58A mice. Hence the mechanism of malignancy in these mice is likely a consequence of aberrant self-renewal activity and resistance to apoptosis in hematopoietic progenitor cells.

We hypothesize that the T58 phosphorylation site on MYC normally acts as a switch to regulate or tune gene expression in response to cytokine stimulation (Fig. 6). Activation of the PI-3K/Akt pathway by cytokines inhibits GSK3β, the kinase that phosphorylates T58, causing T58 to be less phosphorylated (Varano et al. 2017). The change in metabolic state in T58A mutant cells is similar to that seen in germinal center B cells in mice lacking GSK3β, (Jellusova et al. 2017). GSK3β normally restricts glucose metabolism and fatty acid biosynthesis and regulates translation and growth, in part by decreasing the abundance of MYC (Jellusova et al. 2017). In contrast, dephosphorylation of T58 by the Phlpp2 phosphatase stabilizes MYC and has been linked to prostate cancer progression (Nowak et al. 2019). Thus, the Myc phosphorylation switch at T58 integrates cell signaling mediated by GSK3β with the MYC transcriptional program. T58A mutation, by abrogating FBW7-mediated MYC degradation, activates this metabolic switch independently of its normal regulation by GSK3β. This makes T58 a regulatory target for proliferating, self-renewing cancer cells as a means of altering their metabolic state in response to increased biosynthetic demand. This shift in the gene expression program results in increased dependency on glucose metabolism and protein synthesis, and a decreased stress response, which together promote hematopoietic precursor cell survival and self-renewal (Simsek et al. 2010; Ito and Suda 2014). In mature LPS stimulated B cells, this appears to change the balance between protein synthesis and degradation. MYC is critical for proliferation and survival of mature B cells (Tesi et al. 2019) and induces both ribosomal proteins and the unfolded protein response (Iritani and Eisenman 1999; Babcock et al. 2013). However, in the context of the T58A mutation this function is skewed towards modest up-regulation of ribosomal protein production, but decreased UPR.

Genomic occupation by T58A mutant MYC was widespread and increased compared to WT MYC at promoters of expressed genes, whether increased, decreased, or unchanged in expression in MYC-T58A cells. This is consistent with increased abundance of MYC protein due to the T58A block to phospho-degron mediated turnover. While increased MYC occupation had little correlation with gene expression, we did observe an increase in RNA pol2 occupation of gene loci with increased expression in MYC-T58A mutant cells. This is consistent with previous studies indicating that a primary effect of acute activation of MYC on RNA Pol2 is recruitment to promoters to activate transcription (de Pretis et al. 2017), Differentially expressed genes were significantly more likely to be located near MYC-bound enhancers. This raises the possibility that T58A mutant MYC might alter longer range promoter-enhancer interactions. Recent studies indicate that MYC controls chromatin structure in B cells to influence transcription and cell state (Kieffer-Kwon et al. 2017; He et al. 2021; See et al. 2022). For example, a more stable form of MYC may have an increased dwell time on both enhancers and target promoters resulting in augmented transcription of enhancer proximal genes (Fig. 6). Conversely, MYC binding may lead to downregulation of some genes through enhancer interference or a dependence on relatively weaker enhancers. Enhancers can act to negatively regulate gene expression by forming loops into inactive regions of chromatin (Guo et al. 2018). The switch between phosphorylated-unphosphorylated MYC could therefore act to shift 3D chromatin interactions to alter metabolism and survival. Additional studies focused on higher order chromatin structure and direct enhancer-promoter interactions will be required to better understand the regulatory landscape imposed by the T58A mutation.

MYC has also been reported to mediate transcriptional repression through both direct and indirect mechanisms (Kaur and Cole 2013) (Walz et al. 2014). However, the loci showing reduced expression in MYC-T58A cells vs. WT-MYC stimulated B cells (such as pro-apoptotic Bcl2l11, and UPR stress-associated Xbp1) are normally activated by WT-MYC (Muthalagu et al. 2014; de Pretis et al. 2017). Our analysis suggest that these genes are not actively repressed in MYC-T58A cells compared to normal cells, but simply not as strongly activated as in WT MYC cells. Interestingly, we find that many genes exhibiting decreased expression in MYC-T58A mutant cells, relative to WT MYC, are enriched for association with transcription factors regulated by Fbw7 (such as NF-kB/Rel, PU.1, c-JUN, and GFI1), suggesting that the transcriptional effects caused by MYC T58A mutation may result in perturbation of the Fbw7 degradation network. It is noteworthy that in cells with inactive Fbw7, widespread transcriptional changes correlate with regions of open chromatin (Thirimanne et al. 2022).

MYC is part of an extended network of proteins (eg. MNT, MGA, MONDOA) that interact with MAX or the MAX like protein, MLX (reviewed in (Carroll et al. 2018)) and coordinate nutrient availability and cell proliferation. Because MYC-T58A exhibits increased occupancy of promoters and enhancers relative to WT MYC, MAX or other network members may potentially compensate by shifting their genomic binding and patterns of gene expression, as has been reported in studies involving perturbation of the network (Carroll et al. 2021). Experiments designed to address the function of these transcription factors in T58A mutant cells should provide further insight into how this mutation directly and indirectly affects MYC function and its transcriptional program.

## MATERIALS & METHODS

### Generation of T58A mutant mice

To generate a targeting construct replacing the wild-type *Myc* with a *Myc* T58A mutation, a 2.6 kb fragment (Xba1/HindIII digest) of the *Myc* locus (provided by Andreas Trumpp) harboring the first intron and exon 2 of the coding region was subcloned into pBluescript and mutagenized using Pfu DNA polymerase (Stratagene/Agilent) with primers GAGCTGCTTCCCGCACCGCCCCTGTCCC and GGGACAGGGGCGGTGCGGGAAGCAGCTC. This fragment was inserted downstream of a PGKneo cassette flanked by loxP sites in the pLoxP plasmid (provided by Philippe Soriano). An additional 1.2 kb of sequence (HindIII fragment) containing the 3’ UTR of the *Myc* locus was added at the end (for 3’ homology region, HR). At the 3’ end of the HR, a diptheria toxin expression cassette is present (PGK-DTA) to facilitate selection of homologous recombination events. An additional 1.4 kb (BglII/Xba1 fragment) was inserted 5’ of the loxP/PGKneo cassette to provide a 5’ homology arm. The resulting targeting construct was then transfected into ES cells (AK-7), selected with G418, and screened by PCR (using primers specific to the targeting construct and flanking DNA), sequencing, and southern blotting to confirm the mutation and detect proper integration at the *Myc* locus (specify primers and sequence). ES cell clones (two separate clones) exhibiting proper integration were injected into blastocysts and implanted into pseudo-pregnant female mice to generate chimeric pups. Mice containing the T58A allele were then bred with MEOX2-CRE transgenic mouse strain that express germline CRE to excise the intronic PGK-neo cassette in the germline (Tallquist and Soriano 2000). Resulting heterozygous mice harboring the mutated T58A allele were bred together in order to produce mice that were either homozygous for the WT or T58A *Myc* alleles (*Myc^WT/WT^* or *Myc^T58A/T58A^*). Two separately identified ES cell clones were identified that transmitted to the germline and gave reproducible results in early experiments. Determination of mouse genotypes were routinely performed using primers that span the residual loxP site next to the *c-myc* locus (Forward; ACTCGGAGCAGCTGCTAGTC, Reverse: TGCACGTCGCTCTGCTGTTG).

### Mouse strains and breeding

All mouse experiments were approved and performed at the Fred Hutch Cancer Center under IACUC protocols (#1195 and #1450). Myc-T58A mice were backcrossed into C57BL/6 mice for all studies. B6.SJL-Ptprc^a^ Pepc^b^/BoyJ mice (Jackson laboratories), which express the CD45.1 variant of the *Ptprc* gene were used as a source of competitor cells and donor mice for competitive repopulation analysis.

### Western blotting

Tissues and cell pellets were lysed in RIPA buffer. Lysates were quantified by BCA assay (Pierce) or normalized to cell number for equal loading. Samples were resolved on NuPAGE 4-12% Bis-Tris gradient gel before transferring to Nitrocellulose (0.2 micron). Blots were blocked with 5% Milk in TBST, washed with TBST, probed with primary antibody to Myc (Cell Signaling Technologies, D3N8F) or and secondary antibody in 5% Milk in TBST. The secondary antibody was HRP conjugated and chemiluminescent detection was employed. Blots were exposed to Pro-Signal Blotting Film (Genesee Scientific),

### Flow cytometry and cell sorting

For RNA-Seq, CUT&RUN, and ChIP analysis in B cells, single cell suspensions of fresh bone marrow or spleen cells were sorted from *Myc^WT/WT^* or *Myc^T58A/T58A^*mice by positive selection using anti-B220 antibody conjugated MACS beads for marrow derived, immature B cells or depleted for all other markers in mature B cells from spleen (B cell isolation kit, Miltenyi Biotech) using an AutoMACS system (Miltenyi Biotech). Analysis of hematopoietic populations from freshly isolated bone marrow, spleen, and thymus, and sorting of hematopoietic stem (HSC) and progenitor cells (MPP) was carried out using antibodies (FITC, APC or PE conjugated) to differentiated cell markers Mac1/CD11b (Ebioscience, M1/70), Gr1 (Ebioscience, RB6-8C5), F480 (Ebioscience, B6), B220 (Ebioscience, RA3-6B2), CD3e (Ebioscience, 145-2C11), CD4 (BD Bioscience, RM4-5), CD8a (Ebioscience, 53-6.7), and Ter119 (Ebioscience). Analysis and numeration of primitive cells from bone marrow was done using lineage markers, with Sca1/Ly6a (PE or APCcy7 conjugated; Ebioscience, D7), ckit (PE, FITC, or PE-cy7 cojugated; Ebioscience 2B8), and CD150 (Ebioscience, 9D1) antibodies, and proliferating cells were enumerated using AlexaFluor-488 conjugated anti-Ki67 (BD Bioscience, B56) antibody. Apoptosis was detected using FITC labeled VAD-FMK molecule to detect activated caspases (Sigma-Aldrich), or by Annexin V-APC staining (Ebioscience) according to the manufacturer’s protocol. B cells were stained for the glucose analog 2-NBDG (Invitrogen), and Mitotracker red (Invitrogen) as directed by the manufacturer. Intracellular staining using antibodies recognizing MYC (Cell Signaling, D3N8F), Bim (Cell Signaling, C34C5) and Hk2 (Cell Signaling, C64G5) was carried out on methanol fixed cells using intracellular fixation buffer according to the manufacturer (Ebiosciences). For sorting, HSC and progenitor populations prior to competitive repopulation, and single cell analysis, bone marrow cells from *Myc^WT/WT^* or *Myc^T58A/T58A^* mice were first depleted using purified lineage antibodies described above either using anti-rat IgG Dynabeads (ThermoFisher) or MACS lineage depletion beads on an AutoMACS system (Miltenyi Biotech). Cells were then sorted on an Aria3 flow cytometer (Beckton-Dickinson).

### Histology and immunohistochemistry

Mice were euthanized at the time of tumor detection, tissues were collected and fixed in 10% formalin, embedded in paraffin, and stained with Hematoxylin/Eosin (H&E) using standard methodology. For Ki67 immunohistochemistry, fixed tissues (10% formalin) were submitted to the FHCRC histology core for processing. Four-micrometer sections were cut and deparaffinized, followed by heat induced an,gen retrieval. A>er blocking, Ki-67 was detected using a rat an,-Ki67 monoclonal an,body (Dako M7249) at a 1:25 dilu,on followed by bio,nylated goat an,-rat an,body secondary (Jackson ImmunoResearch 112-065-167) and streptavidin-HRP (Jackson ImmunoResearch 016-030-084). Peripheral blood smears from mice with hematopoie,c tumors were stained using Romanowski stain (Diffquick, Fisher). Pictures were taken using a Nikon E800 microscope.

### Competitive repopulation

For competitive repopulation analysis, bone marrow cells were isolated from 3 WT or Myc-T58A mice, and then pooled and sorted for HSCs and MPPs (see Flow cytometry and cell sorting method). Purified HSCs (4-100 cells) and MPPs (40-1000 cells) were then mixed with 1×10^6^ competitor bone marrow cells isolated from B6.SJL-Ptprc^a^ Pepc^b^/BoyJ mice. Cells were then transplanted via tail vein injection into lethally irradiated recipient mice (B6.SJL-Ptprc^a^ Pepc^b^/BoyJ, 50 cGy). Multi-lineage hematopoietic reconstitution was then monitored by obtaining peripheral blood every 3 weeks from recipient mice followed by staining and flow cytometry analysis for differentiated cell markers used above (Mac1, Gr1, B220, CD19, CD4, CD8, F480).

### Culture of hematopoietic progenitors and lymphocytes

For culture of hematopoietic progenitor cells in methylcellulose media, unsorted bone marrow cells from *Myc^+/+^* or *Myc^T58A/T58A^*mice (25,000 cells/ml) were cultured in IMDM containing stem cell factor (SCF Peprotech, 50 ng/ml), IL-6 (Peprotech, 10 ng/ml), IL-3 (Biovision, 10 ng/ml), GM-CSF (Peprotech, 1 ng/ml), 1×10^-4^ M 2-mercaptoethanol, penicillin-streptomycin (10 U/ml penicillin, 10 ug/ml streptomycin), 0.15% methylcellulose (MethoCult M3134, Stem Cell Technologies), and 30% FBS. Hematopoietic progenitor colonies were scored at 7-10 days after culture initiation. For replating experiments, single colonies were picked from the primary cultures, re-cultured in the same media, and scored 7-10 days later. Sorted hematopoietic stem and progenitor cells from Myc^+/+^ or Myc^T58A/T58A^ (500 LSK sorted cells) were cultured in Delta ligand to activate the Notch pathway. For this, wells of 48 well plates were coated with Deltaext-IgG ligand (5 ug/ml), and cultured in IMDM with 100 ng/ml IL-6, SCF, IL-11, and Flt-3 ligand (Biovision). These cultures were maintained by changing media every day. In some experiments, apoptosis was measured by incubating 1 million cells for 1 hour in 1 microliter/ml FITC-VAD-FMK (EMD-Millipore) to detect apoptotic cells by flow cytometry. For lymphocyte cultures, sorted bone marrow or spleen derived B cells were cultured in RPMI with 15% FBS, penicillin-streptomycin, 1×10^-4^ M 2-mercaptoethanol, and either 20 ng/ml IL7 (Biovision) for marrow derived B cells, or LPS (1ug/ml from Sigma-Aldrich) for mature, spleen derived B cells.

### Single cell analysis

Single cell analysis on was carried out on freshly isolated bone marrow cells from either Myc^+/+^ or Myc^T58A/T58A^ 8 week old mice using the 10x Genomics system. Two littermate mice per genotype were sacrificed in two separate experiments (for a total of 4 mice per genotype) and sorted as described above. Cells were then counted and processed according to the manufacturer’s protocol, except that we used a lysis solution with 0.005% digitonin, instead of the suggested 0.01%, in a 2-minute incubation since this gave intact, higher quality nuclei needed for this experiment. Nuclei were then counted and barcoded, followed by sequencing.

After sequencing, alignment was carried out using Cellranger-arc for alignment of multi-ome processed samples (10X genomics). The Cellranger-arc output was then read into the R package Seurat (Stuart et al. 2019), and filtered for high quality, single cells with (RNA amounts between 1000-25,000 total reads per cell) and ATAC less than 70,000 reads per cell). Data was then processed essentially according to the vignette for integration of independent datasets to identify differentially expressed genes (https://satijalab.org/seurat/articles/integration_introduction). After integration and dimension reduction, categories of stem and progenitor cells were identified. Categorization and labelling of each stem and progenitor category was done by comparing to other recent scRNA-Seq datasets (Rodriguez-Fraticelli et al. 2018; Herault et al. 2021) (Karamitros et al. 2018) (Konturek-Ciesla et al. 2023) and identifying marks are show in Supplemental Figure 3A. Most of the more differentiated cell categories were easily defined by expression markers. HSC’s were identifiable by expression of primitive markers such as Meis1 (Nestorowa et al. 2016), and a myeloid/megakaryocyte primed HSC was clearly distinguished by expression of Pbx1 and Vwf (Rodriguez-Fraticelli et al. 2018). Lymphoid progenitors were marked by high levels of Dntt, Flt3, and IL7R. One population expressed primitive markers, with some expression of Flt3, hence was likely a lymphoid primed progenitor, and called MPP1LPrimed (reference (Belluschi et al. 2018; Konturek-Ciesla et al. 2023). The MPP2 population was mainly characterized by increased expression of proliferation markers (Pol1a, Rad51b) as previously characterized (Cabezas-Wallscheid et al. 2014), and called “MPP2Prolif”. Another progenitor cluster characterized by expression of mitotic markers (Kif20b, and Mki67) with relatively high expression of primitive markers was similar to single cell clusters found previously (Konturek-Ciesla et al. 2023), and we called this population “MPP3MPrimed”. Another population of cells expressed both myeloid (Mpo, Irf8) and lymphoid populations (Dntt, Flt3, IL7R) which resembles LMPPs (Karamitros et al. 2018), and we called this LymphoMyeloid.

### RNA-Seq analysis

RNA was extracted with Trizol reagent and quantified on an Agilent 4200 TapeStation. 500ng of RNA was submitted for library preparation through FHCRC Genomics Core. After barcoding and sequencing, libraries were aligned to mm10 using TopHat with then processed using the R package DESeq2. Further analysis was performed using R scripts for plotting in ggplot2 (volcano plots, heatmaps), and enrichment analysis using enrichR, and correlations with other datasets (see Computational Methods).

### ChIP-Seq and Cut&Run

Genomic binding sites for MYC, H3K27Ac, and H3K4Me2 were determined using Cleavage Under Targets and Release Using Nuclease (CUT&RUN) automated using the Auto CUT&RUN system at the FHCRC (Janssens et al. 2018; Skene et al. 2018). Briefly, primary mouse B cells isolated from bone marrow (for Pre B cells), or spleens (for mature B cells) were prepared fresh or stimulated *ex vivo* with IL7 or LPS respectively. Cells were bound to ConA beads, then permeabilized, and incubated overnight with antibody to either c-Myc (Cell Signaling Technologies, D3N8F), H3K27Ac (Abcam, 4729), or H3K4Me2 (EMB Millipore, 07030). After digestion with secondary bound protein-A MNase (Henikoff lab, FHCRC), the reaction was quenched, and 2 ng of yeast ‘spike-in’ DNA was added. DNA fragments released into the supernatant were collected and directly added without purification into an end repair and dA tailing reaction done at 58° (to enhance the capture of small DNA fragments). TruSeq adapters were ligated (Rapid DNA ligase, Enzymatics) onto the DNA, and DNA was digested with Proteinase K. DNA was then size fractionated using Ampure Beads (Beckman-Coulter) and library preparation was then carried out (KAPA Biosystems) by adding barcoded primers. The size distribution was determined for each sample using an Agilent 4200 TapeStation, and quantity of DNA was determined using Qubit (Invitrogen).

Chromatin immunoprecipitation and sequencing (ChIP-seq) was done for RNA-Pol2 (which is poorly detected in Cut&Run) using an MNase digestion step to allow nucleosome resolution of ChIP fragments (Skene and Henikoff 2015). Briefly, after formaldehyde cross-linking, cell lysis, and chromatin fragmentation with MNase, the final SDS concentration after dilution of total chromatin was increased to 0.25% with addition of 20% SDS stock solution. Sonication was performed in a Covaris M220 focused ultrasonicator for 12 minutes with the following settings: 10% duty cycle, 75W peak incident power, 200 cycles/burst, and 6-7°C bath temperature. The SDS concentration of the sonicated chromatin solution was readjusted to 0.1% with dilution buffer. Immunoprecipitation was performed for RNA Pol2 (Active Motif, 4H8) or negative control IgG (Cell Signaling Technology) using 10 μg of antibody for each immunoprecipitation on the clarified chromatin (input) fraction from 10×10^6 cellular equivalents. DNA was then purified using standard phenol:chloroform extraction, 10 pg of spike-in DNA purified from MNase-digested chromatin from *S. cerevisiae* was added to permit comparison between samples. Single strand library preparation for ChIP samples was then performed as described (Ramani et al. 2019).

Samples were then submitted for 25×25 paired-end sequencing (5-10 million reads for CUT&RUN, and 20-30 million reads for ChIP) on an Illumina HiSeq 2500 instrument at the Fred Hutchinson Cancer Research Center Genomics Shared Resource. Sequences were aligned to the mm10 reference genome assembly using Bowtie2 with the arguments: --end-to-end --very-sensitive --no-mixed --no-discordant --overlap --dovetail - I 10 -X 700. Datasets were also aligned to the sc3 *(S. cerevisiae*) assemblies to enumerate reads from spike-in DNA.

Counts per base-pair were normalized as previously described by multiplying the fraction of mapped reads spanning each position in the genome by the genome size or by scaling to spike-in DNA. Peak calling was done using the MACS2 package. For H3K27Ac and H3K4Me2 CUT&RUN data, peaks were called in broad peak mode. Peaks were considered to be associated with a gene if the peak was present within 5kb of the transcription start site (TSS) or in the gene body. Promoters were defined as the −30:300 base position relative the TSS.

### Computational analysis

Downstream analysis of differentially expressed genes by RNA-Seq and correlation with Myc, RNA pol2 occupation, and chromatin modifications with CUT&RUN, ChIP-Seq, and single cell multi-ome data was carried out using custom R scripts and functions developed in the Eisenman lab. Myc, H3K27Ac, and H3K4Me2 occupation for enhancer and promoter occupation representing both Myc^+/+^ or Myc^T58A/T58A^ was determined by merging all files together and calling peaks, and this is how enhancers, and gene bound peaks were identified. The GenomicRanges R package was used to process genome positions. Genomic plots were made using ngs.plot (https://github.com/shenlab-sinai/ngsplot) or the R package ggplot2. Bedtools was also used to numerate genomic positions.

## ACKNOWLEDGEMENTS

We are grateful to Pei-Feng Chang for expert assistance in generating the mutant mice and to Michelle Ulrich for maintaining mouse colonies. We thank David Flowers and Cyd McKay for their valuable help with fluorescent cell sorting, Andreas Trumpp and Philippe Soriano for plasmids, and Xiaoying Wu, Yulong Su, and Markus Welcker for critical readings of the manuscript. We also acknowledge the Fred Hutch Genomics and Bioinformatics, and Comparative Medicine Shared resources for their excellent technical support. Scientific computing infrastructure was supported by ORIP (S10OD028685). This work was supported by NIH NCI grants R35 CA231989 (to RNE), P01HL084205 (to IB), and NIH DP2-HG012442 (to VR).

## AUTHOR CONTRIBUTIONS

Brian Freie, Conceptualization, investigation, formal analysis, writing-original draft; Patrick Carroll: investigation, review and editing; Barbara J. Varnum-Finney, investigation; Vijay Ramani, investigation; Irwin Bernstein; conceptualization, funding acquisition; Robert N Eisenman, conceptualization, supervision, funding acquisition, writing, review and editing.

